# MetaMeta: Integrating metagenome analysis tools to improve taxonomic profiling

**DOI:** 10.1101/138578

**Authors:** Vitor C Piro, Marcel Matschkowski, Bernhard Y Renard

**Affiliations:** Research Group Bioinformatics (MF1) Robert Koch Institute, Nordufer 20, 13353, Berlin, Germany; CAPES Foundation Ministry of Education of Brazil, 70040-020, Brasilia, DF, Brazil

## Abstract

**Background:** Many metagenome analysis tools are presently available to classify sequences and profile environmental samples. In particular, taxonomic profiling and binning methods are commonly used for such tasks. Tools available among these two categories make use of several techniques, e.g. read mapping, k-mer alignment, and composition analysis. Variations on the construction of the corresponding reference sequence databases are also common. In addition, different tools provide good results in different datasets and configurations. All this variation creates a complicated scenario to researchers to decide which methods to use. Installation, configuration and execution can also be difficult especially when dealing with multiple datasets and tools.

**Results:** We propose MetaMeta: a pipeline to execute and integrate results from metagenome analysis tools. MetaMeta provides an easy workflow to run multiple tools with multiple samples, producing a single enhanced output profile for each sample. MetaMeta includes a database generation, pre-processing, execution, and integration steps, allowing easy execution and parallelization. The integration relies on the co-occurrence of organisms from different methods as the main feature to improve community profiling while accounting for differences in their databases.

**Conclusions:** In a controlled case with simulated and real data we show that the integrated profiles of MetaMeta overcome the best single profile. Using the same input data, it provides more sensitive and reliable results with the presence of each organism being supported by several methods. MetaMeta uses Snakemake and has six pre-configured tools, all available at BioConda channel for easy installation (conda install-c bioconda metameta). The MetaMeta pipeline is open-source and can be downloaded at: https://github.com/pirovc/metameta

## Background

A large and increasing number of metagenome analysis tools are presently available aiming to characterize environmental samples [1, 2, 3, 4]. Motivated by the large amounts of data produced from whole metagenome shotgun (WMS) sequencing technologies, profiling of metagenomes has become more accessible and applicable in real scenarios and tends to become the standard method for cheap and fast analysis [5, 6, 7]. Tools which perform sequence classification based on WMS sequencing data come in different flavors. One basic approach is the *de novo* sequence assembly [8, 9, 10], which aims to reconstruct complete or near complete genomes from fragmented short sequences without any reference or prior knowledge. It is the method which provides the best resolution to assess the community composition. However it is very difficult to produce meaningful assemblies from metagenomics data due to short read length, insufficient coverage, similar DNA sequences, and low abundant strains [11].

More commonly, methods use the WMS reads directly without assembly and are in general reference-based, meaning that they rely on previously obtained genome sequences to perform their analysis. In this category of applications, two standard definitions are employed: taxonomic profiling and binning tools. Profilers aim to analyze WMS sequences as a whole, predicting organisms and their relative abundances based on a given set of reference sequences. Binning tools aim to classify each sequence in a given sample individually, linking each one of them to the most probable organism of the reference set. Regardless of their conceptual differences, both groups of tools could be used to characterize microbial communities. Yet binning tools produce individual classification for each sequence and should be converted and normalized to be used as a taxonomic profiler.

Methods available among these two categories make use of several techniques, e.g. read mapping, k-mer alignment, and composition analysis. Variations on the construction of the reference databases, e.g. complete genome sequences, marker genes, protein sequences, are also common. Many of those techniques were developed to overcome the computational cost of dealing with the high throughput of modern sequencing technologies as well as the large number of reference genome sequences available.

The availability of several options for tools, parameters, databases and techniques create a complicated scenario to researchers to decide which methods to use. Different tools provide good results in different scenarios, being more or less precise or sensitive in multiple configurations. It is hard to rely on their output for every study or sample variation. In addition when more than one method is used, inconsistent results between tools using different reference sets are difficult to be integrated. Furthermore, installation, parameterization, database creation as well as the lack of standard outputs are challenges not easily overcome.

We propose MetaMeta, a new pipeline for the joint execution and integration of metagenomic sequence classification tools. MetaMeta has several strengths: easy installation and set-up, support for multiple samples, improved final profile combining multiple tools, out-of-the-box parallelization and high performance computing (HPC) integration, automated databases download and set-up, integrated pre-processing step (read trimming, error correction, and sub-sampling) as well as standardized rules for integration of new tools. MetaMeta achieves more sensitive profiling results than single tools alone by merging their correct identifications and properly filtering out false identifications. MetaMeta was built with SnakeMake [12] and is open-source. The pipeline has six preconfigured tools that can be easily installed through the BioConda channel (https://bioconda.github.io). We encourage the integration of new tools, making it available to the community through a participative Git repository (via pull request). MetaMeta source-code is available at: https://github.com/pirovc/metameta

## Implementation

MetaMeta executes and integrates metagenomic sequence classification tools. The integration is based on several tools’ output profiles and aims to improve organism identification and quantification. An optional pre-processing and sub-sampling step is included. The pipeline is generalized for binning and profiling tools, categories that were previously described in the CAMI (Critical Assessment of Metagenome Interpretation) challenge (http://www.camichallenge.org). MetaMeta provides a pre-defined set of standardized rules to facilitate the integration of tools, easy parallelization and execution in high performance computing infrastructure. The pre-configured tools are available at the BioConda channel to facilitate download and installation, avoiding set-up problems and broken dependencies. The pipeline accepts one or multiple WMS samples and the output is an integrated taxonomic profile for each sample (as well as a separated output from each executed tool). The MetaMeta pipeline can be described in 4 modules: database generation, pre-processing, tool execution, and integration (Figure 1).

**Figure 1:**
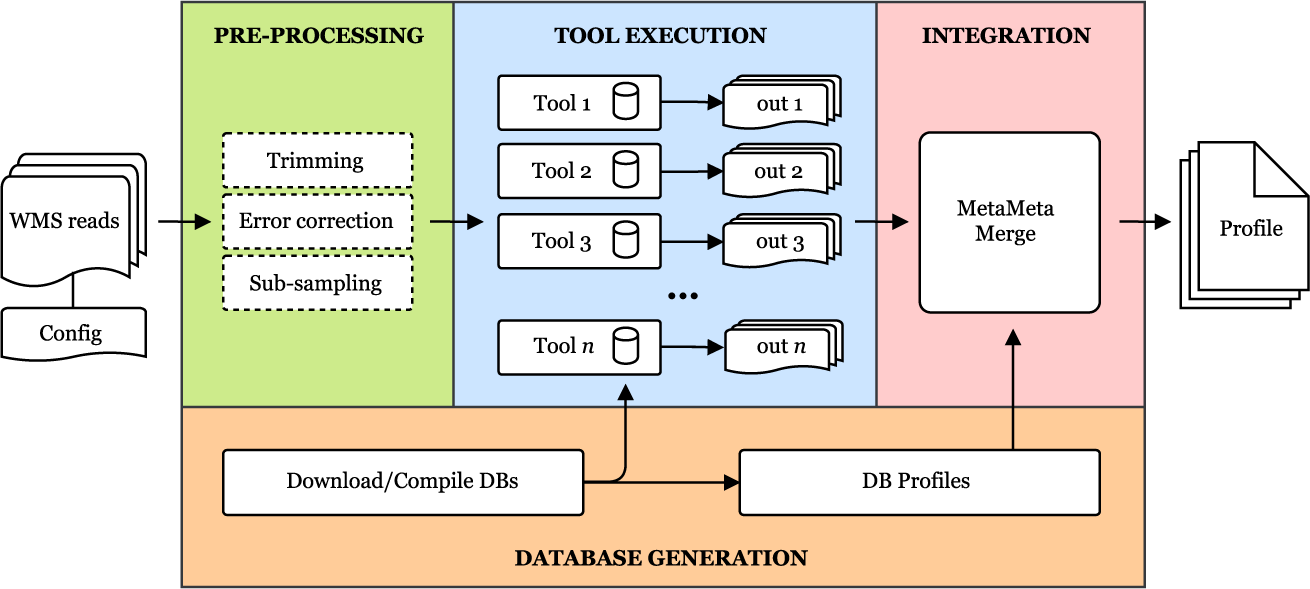
MetaMeta Pipeline. The MetaMeta Pipeline: one or more WMS read samples and a configuration file are the input. The pipeline consists of 4 main modules: Database Generation (only on the first run), Pre-processing (optional), Tool Execution and Integration. The output is a unified taxonomic profile integrating the results from all configured tools for each sample, generated by the MetaMetaMerge module.

## Database generation

On the first run, the pipeline downloads and builds the databases for each of the configured tools. Since each tool has its own database with a specific version of reference sequences, database profiles are generated, collecting which taxonomic groups each tool can identify. Given a list of accession version identifiers for each sequence on the reference set, MetaMeta automatically generates a taxonomic profile for each tool’s database.

## Pre-processing

An optional pre-processing step is provided to remove errors and improve sequence classification: Trimommatic [13] for read trimming and BayesHammer [14] for error correction. A sub-sampling step is also included, allowing the sub division of large read sets among several tools by equally dividing them or by taking smaller random samples with or without replacement, to reduce overall run-time.

## Tool execution

In this step, the pre-processed reads are analyzed by the configured tools. Tools can be added to the pipeline if they follow a minimum set of requirements. They should output their results based on the NCBI Taxonomy database [15] (by name or taxonomic id). Profiling tools should output a rank separated taxonomic profile with relative abundances while binning tools should provide an output with sequence id, length used in the assignment and taxon (More details are given in the Additional file 1). The BioBoxes [16] data format for binning and profiling (https://github.com/bioboxes/rfc/tree/master/data-format) is directly accepted. Tools which provide non-standard output should be configured with an additional step converting their output to be correctly integrated into the pipeline.

## Integration

The integration step will merge identified taxonomic groups and abundances and provide a unified profile for each sample. MetaMeta aims to improve the final results based on the assumption that the more identifications of the same taxon by different tools are reported, the higher its chance to be correct. This task is performed by the MetaMetaMerge module. This module accepts binning and profiling results and relies on previously generated database profiles. Taxonomic classification can change over time and each tool can use a different version/definition of it. For that reason a recent taxonomy database version is used to solve name and rank conflicts (e.g. changing name specification, species turning into sub-species, etc.).

## Abundance estimation binning tools

Binning tools provide a single classification for each sequence in the dataset instead of relative abundances for taxons. An abundance estimation step is necessary for a correct interpretation of such data and posterior integration. The lengths of the binned sequences are summed up for each identified taxonomic group and normalized by the length of their respective reference sequences, estimating the *abundance* for each identified taxon *n* as:

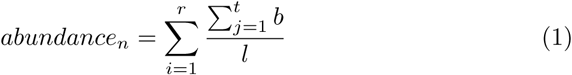

where *r* is the number of reference sequences belonging to the taxonomic group *n, t*_*i*_ is the total of reads classified to the reference *i, b* is the number of bases used in the assignment and *l*_*i*_ is the length of the reference *i*. The abundance of the parent nodes are based on the cumulative sum of their children nodes’ abundance.

## Merging approach

The first step on the merging approach is to normalize estimated abundances to 100% for each taxonomic level. That is necessary because some tools do account for the unclassified reads and others do not. Once normalized, all profiles are then integrated to a single profile. In this step, MetaMetaMerge saves the number of occurrences of each taxon among all profiles. This occurrence count is used to better select taxons that are more often identified, assuming that they have higher chances of being a correct identification. MetaMetaMerge also calculates an integrated value for the relative abundance estimation, defined as the harmonic mean of all normalized abundances for each taxon, avoiding outliers and obtaining a general trend among the estimated abundances. All steps taken in the merging process are performed for each taxonomic level independently, from super kingdom to species by default.

Since tools use different databases of reference sequences it is necessary to account for this bias. Previously generated database profiles provide which taxons are available for each tool. By merging all database profiles, it is possible to anticipate how many times each taxon could be identified among all tools used. The number of occurrences of each taxon from the tools’ output and the database presence number are integrated to generate a score *S* for each taxon, defined as:

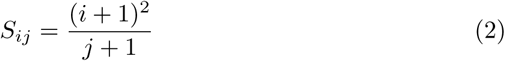

where *i* is the number of times the taxon was identified and *j* the number of times it is contained in the databases. This score calculation accounts for the presence/absence of taxonomic groups on different databases. It gives higher scores to the most identified taxons present in more databases. At the same time, lower scores are assigned to taxons present in many databases but not identified too many times. The score calculation is purposely biased for higher scores when *i* = *j* (Additional file 1: Figure 1), given the benefit of the doubt for taxons with low identification that are available only in few databases.

Commonly, metagenome analysis methods have to deal with a moderate to high number of false positive identifications at lower taxonomic levels. That occurs mainly because metagenomes can contain very low abundant organisms with similar genome sequences. This problem is even extended in our merged profile by collecting all false positives from different methods, generating a long tail of false positives with lower scores mixed together with true identifications. A filtering step is therefore necessary to avoid wrong assignments. This step is usually performed by an abundance cutoff value. Setting up this value is subject to uncertainty since the real abundances are usually not known and the separation between low abundant organisms and false identifications is not clear [17]. A simple cutoff would not provide a good separation between true and false results in this scenario.

To overcome this problem, MetaMetaMerge classifies each taxon in a set of bins (four by default) based on the calculated score (Equation 2). Bins are defined by equally dividing the range of scores in the numbers of bins selected. Now each taxon has a score and a bin assigned to it. Taxons with higher scores are more likely to be true identifications and are going to be grouped together in the same bin. With this strategy it is possible to obtain a general separation among taxons which are prone to be true or false identifications. Within each taxon grouped in a bin (sorted by relative abundance) a cutoff is applied to remove possible false identifications with low abundance. Here, the cutoff value can be selected based on pre-defined functions, which can achieve more sensitive or precise results (Additional file 1: Mode functions). After the cutoff, the remaining taxons are grouped together and sorted by relative abundance to generate the final profile.

At the end, MetaMeta will provide a final taxonomic profile, integrating all tools results, a detailed profile with co-occurrence and individual abundances as well as single profiles for each executed tool.

## Results

### Tool selection

MetaMeta was evaluated with a set of six tools: CLARK [18], DUDes [19], GOTTCHA [20], Kraken [21], Kaiju [22], and mOTUs [23]. The choice was motivated by recent publications comparing the performance of such tools [3, 4]. We also selected new tools that represent state-of-the-art of sequence classification for metagenomics data, as long as they fit in our pipeline requirements described on the Methods section. We considered the amount of data/run time performance for each tool, selecting only the ones that can handle large amounts of data as commonly used today in metagenomics analysis in an acceptable time (for our largest dataset less than 1 day). We also selected an equal number of tools for each category: DUDes, GOTTCHA and mOTUs are taxonomic profiling tools, while CLARK, Kraken and Kaiju are binning tools. Databases were created following the default guidelines for each tool, considering only bacteria and archaea as targets.

### Datasets and evaluation

The pipeline was evaluated with a set of simulated and real samples (Table 1). The simulated data were provided as part of the CAMI Challenge (toy samples) and the real samples were obtained from the Human Microbiome Project (HMP) [24, 25]. MetaMeta was compared to each single result from each tool configured in the pipeline. Although the pipeline can work on the strain level, we evaluate the results until species levels since most of the tools still do not provide strain level identifications. We compare the results to the ground truth in a binary (true and false positives, sensitivity and precision) and quantitative way (*L*_1_ norm) if abundance profiles were available. Computer specifications and parameters can be found on the Additional file 1.

**Table 1:** Samples used in this study. * expected number of species from isolated genomes from the gastrointestinal tract

| Sets | # Samples | Total bases | # species |
| --- | --- | --- | --- |
| CAMI Toy Low | 1 | 14.8 Gbp | 30 |
| CAMI Toy Medium | 4 | 31.3 Gbp | 199 |
| CAMI Toy High | 5 | 74.5 Gbp | 375 |
| HMP Stool | 10 | 102.2 Gbp | 299* |

### CAMI data

The CAMI challenge provided three toy datasets of different complexity (Table 1) with known composition and abundances. From low to high complexity, they provide an increasing number of organisms and samples. The samples within a complexity group contain the same organisms with variable abundances among samples. The sets contain real and simulated strains from complete and draft bacterial and archaeal genome sequences. The simulated CAMI datasets, especially those of medium and high complexity, provide a very challenging and realistic data in terms of complexity and size.

In Figure 2 it is possible to observe the tools performance in terms of true and false positives for the CAMI high complexity set. All configured tools perform similarly in the true positive identifications but vary among the false positives. Binning tools have a higher number of false positive identifications due to the fact that even single classified reads are considered. MetaMetaMerge profile surpassed all other methods in true positive identifications while keeping the false positive number low. The same trend occurs in the other complexity sets (Additional file 1: Figure. 3–8). Figure 3 shows the trade-off between precision and sensitivity for all high complexity samples. MetaMetaMerge achieved the best sensitivity while GOTTCHA the best precision among the compared tools with default parameters. Those results show how the merging module of the MetaMeta pipeline is capable of better selecting and identifying true positives based on the co-occurrence information. MetaMetaMerge also has the flexibility to provide more precise or sensitive results (Figure 3) just by changing the *mode* parameter (details are given in the Additional file 1: Mode functions). In the very precise mode, the merged profile outperformed all tools in terms of precision, but with the cost of losing sensitivity. In the very sensitive mode, the merged profile could improve the sensitivity compared to the run with default parameters, with some loss of precision. It is important to notice that the trade-off between precision and sensitivity could also be explored by the *cutoff* parameter (default 0.0001), depending on what is expected to be the lowest abundant organism in the sample. The MetaMetaMerge *mode* parameter will give more precise or sensitive results based on this cutoff value.

**Figure 2:**
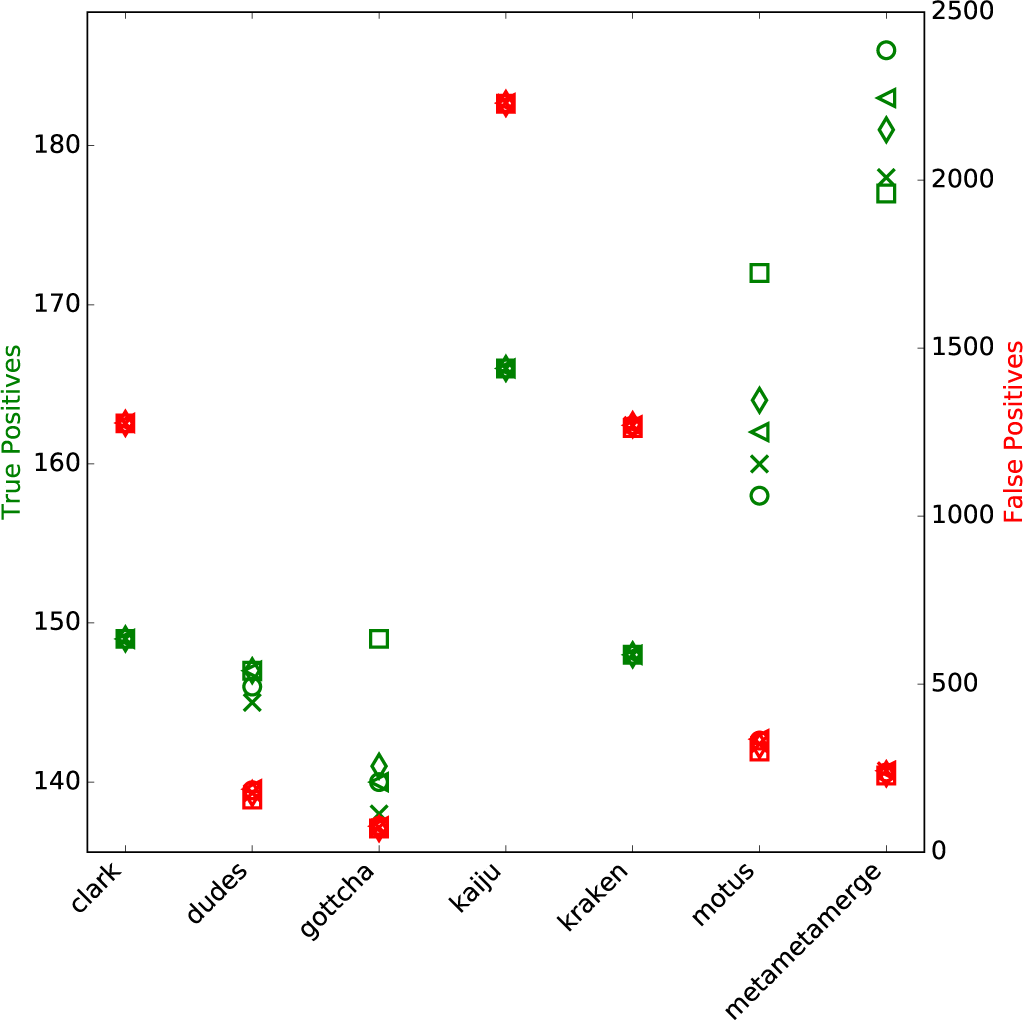
True and False Positives CAMI high complexity set. In green (left y axis): True Positives. In red (right y axis): False Positives. Results at species level. Each marker represents one out of five samples from the CAMI high complexity set.

**Figure 3:**
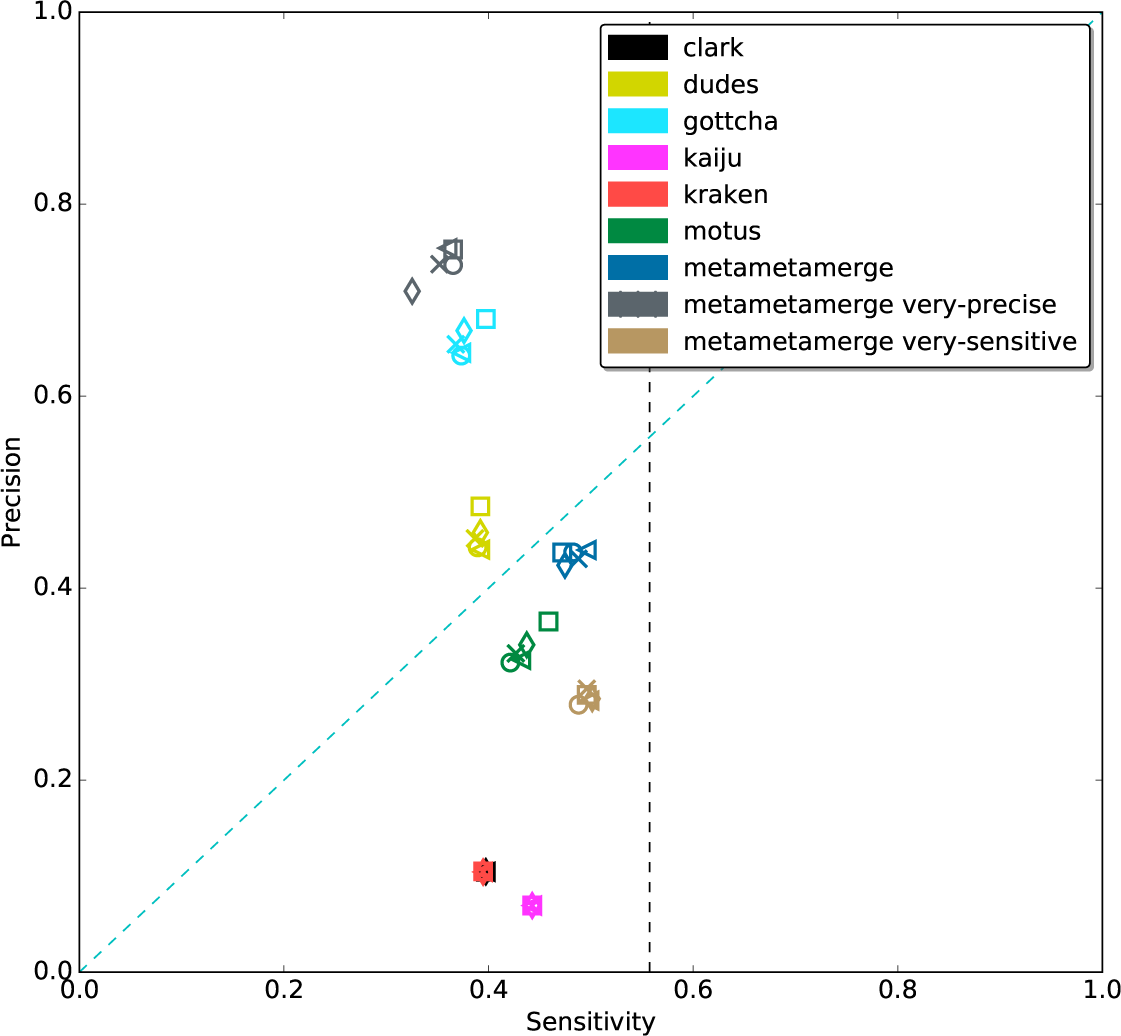
Precision and Sensitivity CAMI high complexity set Dotted black line marks the maximum possible sensitivity value (0.57) that could be achieved with the given t ools and databases. Results at species level. Each marker represents one out of five samples from the CAMI high complexity set.

In terms of relative abundance, MetaMetaMerge provides the most reliable predictions with smaller difference from the real abundances, as shown in Figure 4 with regard to the *L*_1_ norm measure. By taking the harmonic mean, we succeed in reducing the effect of outliers that occur among the tools and capture the trend of the estimated relative abundances, providing a new, more robust estimate.

**Figure 4:**
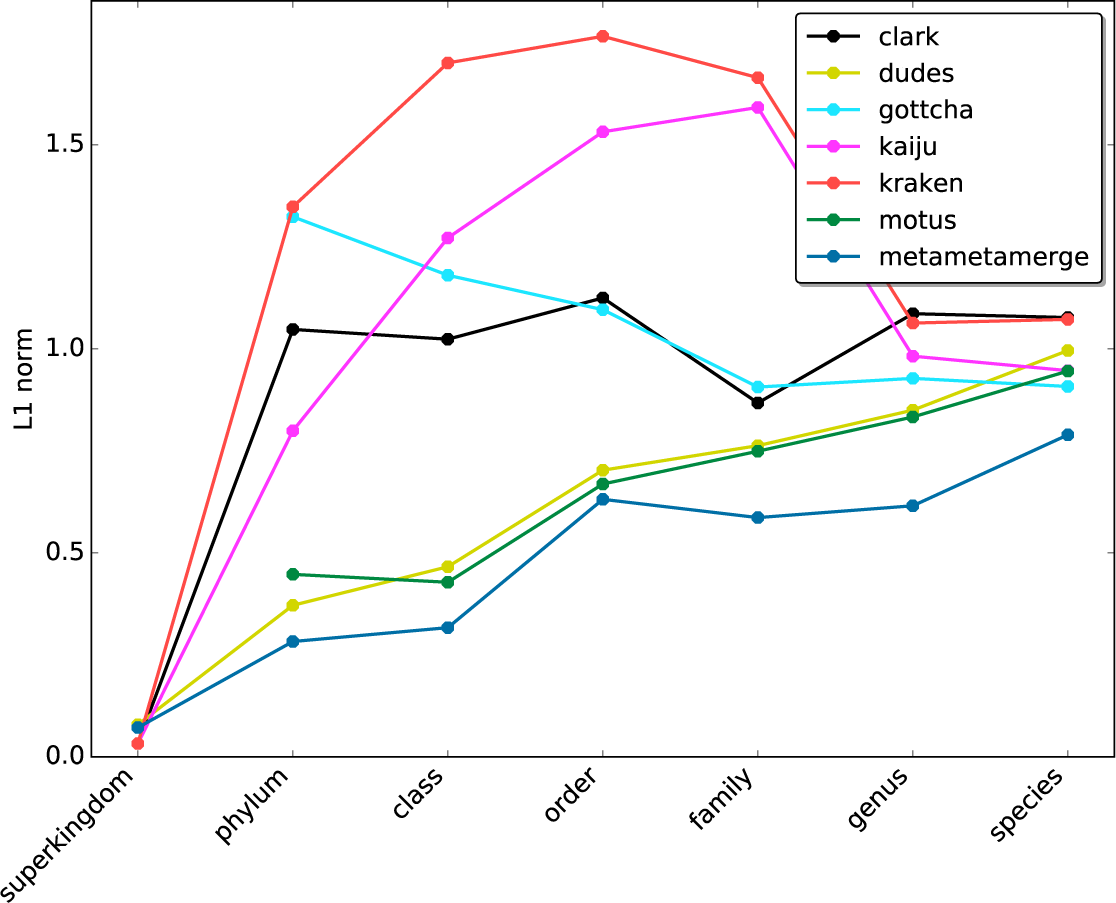
*L*_1_ **norm error** Mean of the *L*_1_ norm measure at each taxonomic level for five samples from the high complexity CAMI set.

### Pre-processing and sub-sampling effects

We explore here the effects of pre-processing and sub-sampling on the CAMI toy sets. Results shown in this section were trimmed and sub-sampled in several sizes, with and without replacement and executed five times for each sub-sample. Trimming effects were small on this set, slightly increasing precision (data not shown). Figure 5 shows the effects of sub-sampling in terms sensitivity and run-time (full pipeline) for one of the high complexity CAMI sets. Sampling provides a high decrease on run-time for every tool and consequently for the whole pipeline. However, only below 5% it is possible to see a significant but still small decrease on sensitivity. All tools behave similarly on the sub-sampled sets, with GOTTCHA and mOTUs having a high decrease of sensitivity when using only 1% of the data. With the same sub-sample configuration (1%), MetaMetaMerge achieved a sensitivity higher than any other tool alone using 100% of the set. It also runs the whole pipeline approximately 17 times faster than with the full set (from 05h41m36s to 20m19s on average), being faster than the fastest tool with 100% of the data (kraken 29m26s on average) and the second best sensitive tool (kaiju 1h47m44 on average). As expected, precision is slightly increased in small sub-samples due to less data (Additional file 1: Figure 9).

**Figure 5:**
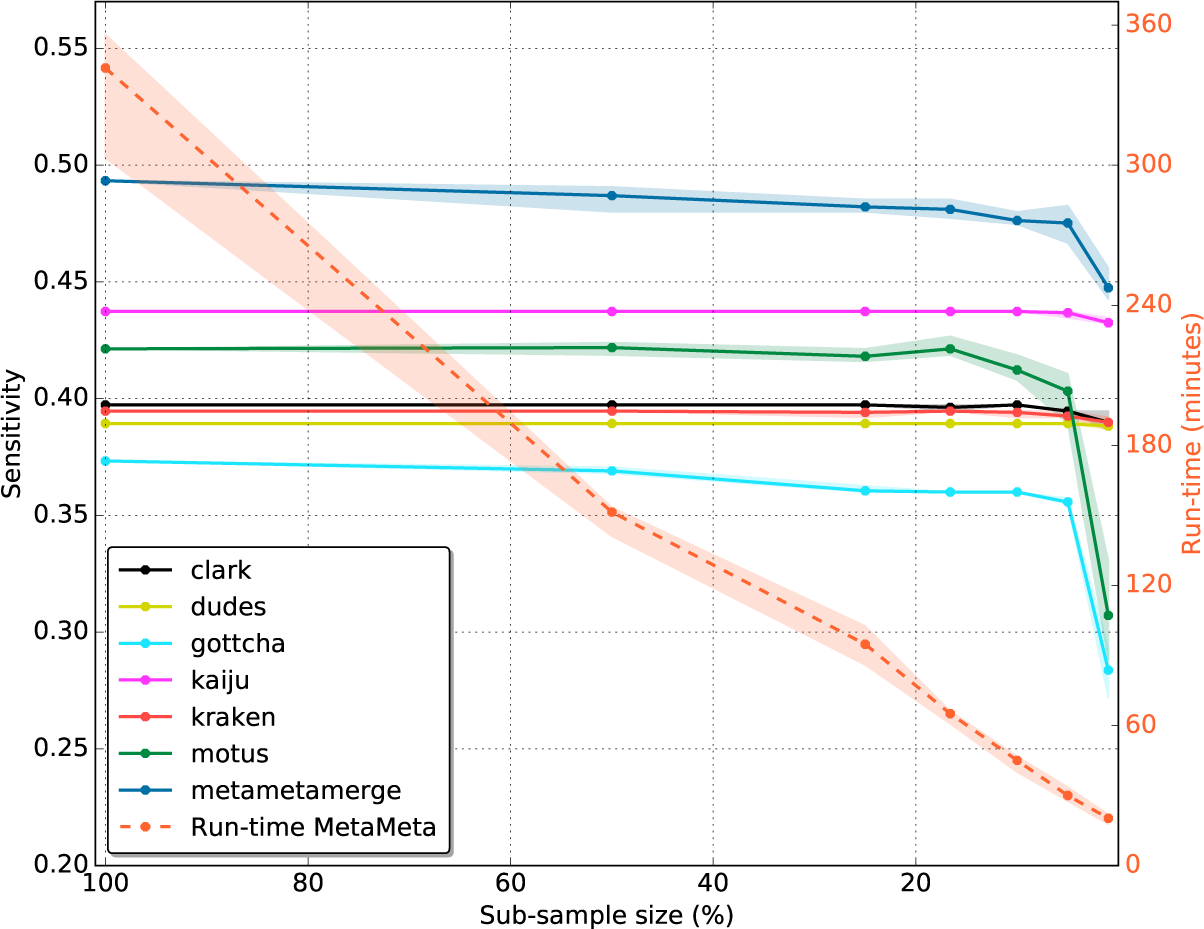
sub-sampling. Sensitivity (left y axis) and run-time (right y axis) at species level for one randomly selected CAMI high complexity sample. Each sub-sample was executed five times. Lines represent the mean and the area around it the maximum and minimum achieved values. Run-time stands for the time to execute the MetaMeta pipeline. The evaluated sample sizes are: 100%, 50%, 25%, 16.6%, 10%, 5%, 1%. 16.6% is the exact division among 6 tools, using the the whole sample. sub-samples above that value were taken with replacement and below without replacement. The plot is limited to a value of 0.57 (left y axis) that is the maximum possible sensitivity value that could be achieved with the given tools and databases.

### Human Microbiome Project data

The HMP provided several resources to characterize the microbial communities at different sites of the human body. MetaMeta was tested on stool samples to evaluate the performance of the pipeline on real data. For evaluation we use a list of reference genome sequences that were isolated from specific body sites and sequenced as part of the HMP. They do not represent the complete content of microbial diversity in each community but serve as a guide to check how well the tools are performing. Randomly selected stool samples were compared against the isolated genomes obtained from the gastrointestinal tract.

Figure 6 shows the results for 10 randomly selected samples. In sensitive mode, MetaMetaMerge achieved the highest number of true positive identifications with a moderate number of false positives, below all binning tools but slightly above the taxonomic profilers. mOTUs produced good results in the selected samples mainly because its database is based on the isolated genomes from the HMP (the same as the ground truth used here). Since mOTUs is the only tool with a distinct set of reference sequences that could classify this set, the scores (from Equation 2) attributed to mOTUs’ unique identifications were low. Still, MetaMetaMerge could improve the true identifications keeping a lower rate of false positives by incorporating the true identifications from other methods.

**Figure 6:**
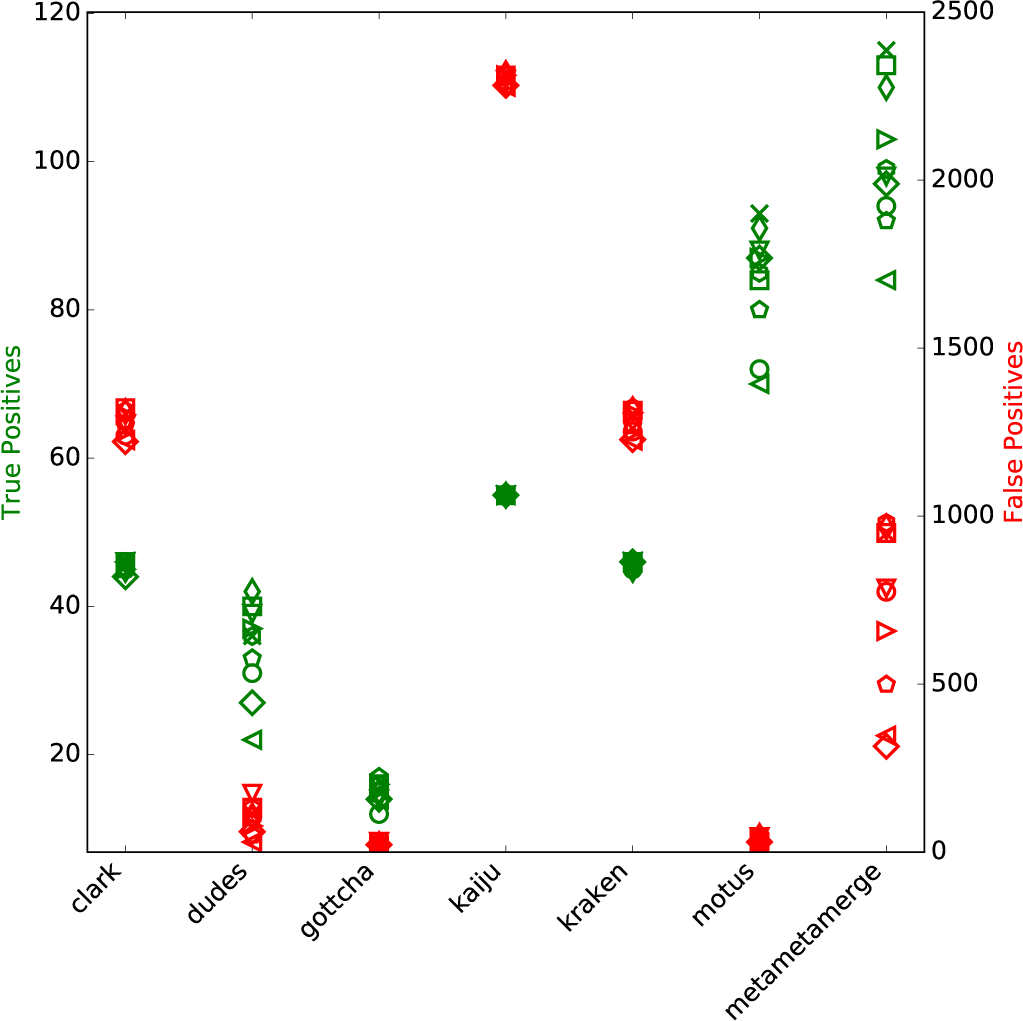
True and False Positives HMP stool samples. In green (left y axis): True Positives. In red (right y axis): False Positives. Results at species level. Each marker represents one out of ten randomly selected stool samples from the HMP.

## Discussion

MetaMeta is a complete pipeline for classification of metagenomic datasets. It provides improved profiles over the tested tools by merging their results. In addition, the pipeline provides easy configuration, execution and parallelization. With simulated and real data, MetaMeta is capable to achieve higher sensitivity. That is possible due to the MetaMetaMerge module, which extracts information of co-occurrence of taxons on databases and profiles. Using this information with a novel approach, MetaMetaMerge avoids false positives and keeps most of the true identifications, exploring the complementarity of currently available methods.

By running several tools, MetaMeta has an apparently prohibitive execution time. In reality, the parallelization provided by Snakemake makes the pipeline run in a reasonable time using most of the computational resources. That is possible by the way the rules are chained and executed among several cores, lasting not more than the slowest tool plus pre- and post-processing time, which are very small in comparison to the analysis time. In addition, sub-sampling allows the reduction of input data and a high decrease of execution time with small if any impact on the final result. That is viable due to redundant data contained in many metagenomic samples as well as redundant execution by several tools provided in the MetaMeta environment. However sub-sampling should be used with caution, taking in consideration the coverage of low abundant organisms.

All tools presented here are available at the BioConda channel and can be installed with a single command, working out-of-the-box for several computer environments and avoiding conflicts and broken dependencies. MetaMeta can also handle multiple large samples at the same time, with options to delete intermediate files and keep only necessary ones, being well suited to large scale projects. It also reduces idle computational time by smartly parallelizing samples among one or more tools (Additional file 1: Figure. 10-12). The parallelization noticeably decreases the run time when computational power is available and manages to serialize and control the run when access to computational power is limited. Integration into HPC systems is also possible and we provide a pre-configured file for queuing systems (e.g. slurm). As stated by Lee et al. [26], solid-state drives accelerate the run time of many bioinformatics tools. Such drives were used in the evaluations shown in this paper and are beneficial for the MetaMeta pipeline.

MetaMeta makes it easier for the user to obtain more precise or sensitive results by providing a single default parameter as well as advanced options for more refined results. All other tools were used in default mode, meaning that it is possible to obtain problem-centric optimized results only by changing the way MetaMeta works. That facilitates and simplifies the task for researchers that are in search for a specific goal.

MetaMeta supports strain level identification. Nevertheless all evaluations were made at species level due to lack of support to strain identification in some tools. Also the lack of standard was a limiting factor. Taxonomic IDs are no longer assigned to strain levels [27] and tools output them in different ways. With standard output definitions, the use of strain classification on the pipeline is straight forward.

Related in parts, a method called WEVOTE was developed in parallel and recently published [28] where five classification tools were used to generate a merged taxonomic profile. Although the two methods present distinct ways of achieving better taxonomic profiling, they are not built for the same use case. WEVOTE relies on BLAST based tools and thereby is not suited for the large scale WMS applications, since the dataset sizes practically prohibit analyses via BLAST based approaches. Differently, MetaMeta was built accounting for high throughput data. Moreover, we supply an easy way to install tools and MetaMeta provides a complete pipeline which can configure databases and run classification tools with an integration module at the end, where WEVOTE provides only the integration method. As a result a comparison among the pipelines is hard to perform and interpret since they both use a different set of tools and databases.

In conclusion, MetaMeta is an easy way to execute and generate improved taxonomic profiles for WMS samples with multiple tool support. We believe the method can be very useful for researchers that are dealing with multiple metagenomic samples and want to standardize their analysis. The MetaMeta pipeline was built in a way to facilitate the execution in many computational environments using Snakemake and BioConda. That diminishes the burden of installing and configuring multiple tools. The pipeline also gives control over the storage of the results and has an easy set of parameters which makes it possible to obtain more precise or sensitive results. MetaMeta was coded in a standardized manner, allowing easy expansion to more tools, also collectively in the MetaMeta git repository (https://github.com/pirovc/metameta). We believe that the final profile could be even further improved with novel tools configured into the pipeline.

## List of abbreviations

CAMI: critical assessment of metagenome interpretation
HMP: human microbiome project
HPC: high performance computing
WMS: whole metagenome shotgun

## Ethics approval and consent to participate

This manuscript does not report data collected from humans or animals

## Consent for publication

This manuscript does not contain any individual person’s data in any form

## Availability of data and material

The software presented in this manuscript is available at: https://github.com/pirovc/metameta/ and https://github.com/pirovc/metametamerge/. CAMI data sets are available at: http://data.cami-challenge.org/. HMP data sets are avaiable at NCBI Sequence Read Archive: https://www.ncbi.nlm.nih.gov/sra with the following accession numbers: SRS013476, SRS014979, SRS015065, SRS016018, SRS016267, SRS016989, SRS017191, SRS022071, SRS049712, SRS065504.

## Competing interests

The authors declare that they have no competing interests.

## Funding

This work was supported by the Coordenação de Aperfei¸coamento de Pessoal de Nível Superior (CAPES) Ciencia sem Fronteiras [BEX 13472/13-5 to VCP] and by the German Federal Ministry of Health [IIA5-2512-FSB-725 to BYR].

## Author’s contributions

VCP and BYR conceived the project and designed the methods. VCP developed the pipeline and MM led the sub-sampling analysis. VCP and BYR interpreted the data. VCP drafted the manuscript with contributions by MM and BYR. All authors read and approved the final manuscript.

## Acknowledgements

We thank Enrico Seiler for proof-reading the manuscript and for technical support.

## Tables Additional Files Additional file 1

Additional File with supplementary figures and information

## References

[1] Adam L Bazinet and Michael P Cummings. A comparative evaluation of sequence classification programs. BMC Bioinformatics, 13(1):92, 2012.

[2] Pavlopoulos, Anastasis Oulas, Christina Pavloudi, Paraskevi Polymenakou, Nikolas Papanikolaou, Georgios Kotoulas, Christos Arvanitidis, and Ioannis Iliopoulos. Metagenomics: Tools and Insights for Analyzing Next-Generation Sequencing Data Derived from Biodiversity Studies. Bioinformatics and Biology Insights, page 75, May 2015.

[3] Michael a. Peabody, Thea Van Rossum, Raymond Lo, and Fiona S. L. Brinkman. Evaluation of shotgun metagenomics sequence classification methods using in silico and in vitro simulated communities. BMC Bioinformatics, 16(1):363, dec 2015.

[4] Stinus Lindgreen, Karen L. Adair, and Paul P. Gardner. An evaluation of the accuracy and speed of metagenome analysis tools. Scientific Reports, 6:19233, jan 2016.

[5] Claudio U K¨oser, Matthew J Ellington, Edward J P Cartwright, Stephen H Gillespie, Nicholas M Brown, Mark Farrington, Matthew T G Holden, Gordon Dougan, Stephen D Bentley, Julian Parkhill, and Sharon J Peacock. Routine use of microbial whole genome sequencing in diagnostic and public health microbiology. PLoS pathogens, 8(8):e1002824, January 2012.

[6] M. J. Pallen. Diagnostic metagenomics: potential applications to bacterial, viral and parasitic infections. Parasitology, 141(14):1856–1862, December 2014.

[7] W Florian Fricke and David a Rasko. Bacterial genome sequencing in the clinic: bioinformatic challenges and solutions. Nature reviews. Genetics, 15(1):49–55, January 2014.

[8] Toshiaki Namiki, Tsuyoshi Hachiya, Hideaki Tanaka, and Yasubumi Sakakibara. MetaVelvet: an extension of Velvet assembler to de novo metagenome assembly from short sequence reads. Nucleic Acids Research, 40(20):e155 – e155, nov 2012.

[9] Yu Peng, Henry C M Leung, S M Yiu, and Francis Y L Chin. IDBA-UD: a de novo assembler for single-cell and metagenomic sequencing data with highly uneven depth. Bioinformatics, 28(11):1420–8, jun 2012.

[10] D. Li, C.-M. Liu, R. Luo, K. Sadakane, and T.-W. Lam. MEGAHIT: An ultra-fast single-node solution for large and complex metagenomics assembly via succinct de Bruijn graph. Bioinformatics, pages btv033–, 2015.

[11] Adina Howe and Patrick S G Chain. Challenges and opportunities in understanding microbial communities with metagenome assembly (accompanied by IPython Notebook tutorial). Frontiers in Microbiology, 6(JUL):10–13, July 2015.

[12] J. Koster and Sven Rahmann. Snakemake–a scalable bioinformatics workflow engine. Bioinformatics, 28(19):2520–2522, October 2012.

[13] Anthony M Bolger, Marc Lohse, and Bjoern Usadel. Trimmomatic: a flexible trimmer for Illumina sequence data. Bioinformatics, 30(15):2114–2120, aug 2014.

[14] Sergey I Nikolenko, Anton I Korobeynikov, and Max a Alekseyev. BayesHammer: Bayesian clustering for error correction in single-cell sequencing. BMC Genomics, 14(Suppl 1):S7, 2013.

[15] Scott Federhen. The NCBI Taxonomy database. Nucleic Acids Research, 40(D1):D136–D143, jan 2012.

[16] Peter Belmann, Johannes Dr¨oge, Andreas Bremges, Alice C. McHardy, Alexander Sczyrba, and Michael D. Barton. Bioboxes: standardised containers for interchangeable bioinformatics software. GigaScience, 4(1):47, dec 2015.

[17] M. L. Zepeda Mendoza, T. Sicheritz-Ponten, and M. T. P. Gilbert. Environmental genes and genomes: understanding the differences and challenges in the approaches and software for their analyses. Briefings in Bioinformatics, (November 2014):1–14, feb 2015.

[18] Rachid Ounit, Steve Wanamaker, Timothy J Close, and Stefano Lonardi. CLARK: fast and accurate classification of metagenomic and genomic sequences using discriminative k-mers. BMC Genomics, 16(1):236, dec 2015.

[19] Vitor C. Piro, Martin S. Lindner, and Bernhard Y. Renard. DUDes: a topdown taxonomic profiler for metagenomics. Bioinformatics, 32(15):2272–2280, aug 2016.

[20] T. a. K. Freitas, P.-E. Li, M. B. Scholz, and P. S. G. Chain. Accurate read-based metagenome characterization using a hierarchical suite of unique signatures. Nucleic Acids Research, 43(10):e69–e69, may 2015.

[21] Derrick E Wood and Steven L Salzberg. Kraken: ultrafast metagenomic sequence classification using exact alignments. Genome biology, 15(3):R46, mar 2014.

[22] Peter Menzel, Kim Lee Ng, and Anders Krogh. Fast and sensitive taxonomic classification for metagenomics with Kaiju. Nature Communications, 7:11257, apr 2016.

[23] Shinichi Sunagawa, Daniel R Mende, Georg Zeller, Fernando Izquierdo-Carrasco, Simon a Berger, Jens Roat Kultima, Luis Pedro Coelho, Manimozhiyan Arumugam, Julien Tap, Henrik Bjørn Nielsen, Simon Rasmussen, Søren Brunak, Oluf Pedersen, Francisco Guarner, Willem M de Vos, Jun Wang, Junhua Li, Jo¨el Dor´e, S Dusko Ehrlich, Alexandros Stamatakis, and Peer Bork. Metagenomic species profiling using universal phylogenetic marker genes. Nature Methods, 10(12):1196–1199, oct 2013.

[24] Barbara A. et al. Methé. A framework for human microbiome research. Nature, 486(7402):215–221, June 2012.

[25] Curtis et al. Huttenhower. Structure, function and diversity of the healthy human microbiome. Nature, 486(7402):207–214, June 2012.

[26] Sungmin Lee, Hyeyoung Min, and Sungroh Yoon. Will solid-state drives accelerate your bioinformatics? In-depth profiling, performance analysis and beyond. Briefings in Bioinformatics, 17(4):713–727, July 2016.

[27] Scott Federhen, Karen Clark, Tanya Barrett, Helen Parkinson, James Ostell, Yuichi Kodama, Jun Mashima, Yasukazu Nakamura, Guy Cochrane, and Ilene Karsch-Mizrachi. Toward richer metadata for microbial sequences: replacing strain-level NCBI taxonomy taxids with BioProject, BioSample and Assembly records. Standards in Genomic Sciences, 9(3):1275–1277, January 2014.

[28] Ahmed A. Metwally, Yang Dai, Patricia W. Finn, and David L. Perkins. WEVOTE: Weighted Voting Taxonomic Identification Method of Microbial Sequences. PLOS ONE, 11(9):e0163527, September 2016.

